# Theta and beta account for the two-impacted components of social influence on risk decision-making

**DOI:** 10.64898/2025.12.16.694560

**Authors:** Yixuan Lin, Xiaoyu Zhou, Xuena Wang, Yangzhuo Li, Xianchun Li

**Author notes:** **Corresponding author:** School of Psychology and Cognitivse Science, East China, Normal University, Shanghai, 200062, China. **Competing interest statement:** The author(s) declared that there were no conflicts of interest with respect to the authorship or the publication of this article. **Data availability statement:** The data and code have been uploaded to the Open Science Framework (OSF) website: behavioral and modeling data and codes: https://doi.org/10.17605/OSF.IO/UP6GE; EEG data and codes: https://doi.org/10.17605/OSF.IO/XTUWE. **Human ethics statements:** The study was approved by the Human Research Protection Committee of East China Normal University (No. HR 429-2020).

## Abstract

Social influence in risky decision-making means changing risky behavior after observing other’s choices. Although attributes’ prioritization can alter subjective value of a particular option while choice bias can affect the tendency toward it, previous studies have not simultaneously examined these two components. Here, participants made risky or safety choices, observed a model whose risk preference either matched (control) or opposed (inconsistent) their own, then repeated the task. Behavioral and EEG data were collected, and drift-diffusion model (DDM) was used to decompose different components. After observing an opposing preference, risk-seeking participants chose safety options more often, and risk-aversion participants chose risky options more often, indicating social influence. Model comparison favored an attribute-wise DDM. Social influence altered two distinct components in risky decisions: the weight of amount in value computation (more negative after observing the opposing model) and the starting point (shifting toward the options chosen more often). Multivariate pattern analysis linked these components to distinct neural signatures: theta activity (≈500 ms after option onset) tracked the amount weighting, whereas pre-option beta activity tracked the starting point. Together, the results show that social influence reshapes risky decisions by reweighting amount and shifting choice bias, each supported by dissociable electrophysiological dynamics.

## 1 Introduction

Life often presents situations requiring risky decisions, such as financial investments, substance use, or choices about further education and career paths. Individual risk preferences can be shaped by external factors ^1–3^, with social influence playing a particularly important role^4^. For example, after observing others whose risk preferences inconsistent with their own, individuals’ preferences can reverse: risk-seeking individuals may become more risk-averse, and vice versa ^5^. While effect of social influence on risky decision-making has been documented and prior studies have mainly focused on identifying the associated brain regions, how social influence modulates the decision-making process remains unclear.

The comparison of subjective values between options drives individual decision-making. Previous studies have typically integrated amount and probability as a single entity, using an option-wise approach for computational modeling and identifying neural correlates of social influence ^5–7^. However, the impact of social influence on these attributes might differ. Others’ goals can shape one’s preferences and choices ^8,9^. Specifically, individuals can learn about and represent others’ goal through observation ^10^, reshaping the prioritization of different attributes during decision-making and guiding choices to align with the goal ^11^. Recent study has provided supporting evidence that these attributes are compared independently ^12^. Thus, social influence likely alters risky decision-making at the attribute-wise way rather than the option-wise way, yet this has not been empirically examined. Moreover, it was also proposed that probability might be completely ignored when in uncertainty ^13^ and the priority heuristic theory suggested that individuals prioritize amount over probability when evaluating options in risky decision-making ^14^. Therefore, we further hypothesize that social influence may reshape risky decision-making by altering the prioritization of amount in subjective value.

Additionally, previous researches overlooked the potential alterations of social influence on choice bias. During decision-making, individuals can exhibit a predisposition toward a particular option before evaluating different attributes, which is known as choice bias. For instance, individuals were inclined to avoid risky options in decision-making to prevent bad outcomes before considering possible gains or losses ^15^. Generally, individuals would imitate the behavior of others outside of conscious awareness and intent, showing a bias toward merely copying decisions of others ^16,17^. Previous research suggested that the change of observational learning on choice bias might moderated by the type of decision. It was showed that observing an antisocial peer’s moral decisions can increase the bias towards choosing harmful option ^18^. However, when the assignment plan was highly unfair, observing others’ decisions to punish did not change the choice bias towards compensatory or punitive options ^19^. Although value computation and choice bias can lead to the same results in risk decision-making, they shape the decision process in different ways and involve different physiological mechanisms ^20^. Accordingly, the impact of social influence on choice bias in risk decision-making needed to be further investigated.

At the behavioral level, these two distinct components can be examined via drift-diffusion model (DDM). It proposed that decisions were not made instantly, but that choices were ultimately made through a process of evidence accumulation over time ^21,22^. In the DDM, each decision is represented by an upper and a lower boundary. Individuals start their decision process at a point between these boundaries and accumulate evidence over time until cross one of the boundaries, leading to the corresponding response. A recent study has showed that attribute-wise models fit intertemporal decision-making better than option-wise models via DDM ^23^. Thus, the DDM is well-suited for identifying how social influence changes the prioritization of different attributes (parameter: weight in evidence accumulation) and the choice bias (parameter: starting point) in risk decision-making.

Besides computational modeling, electroencephalography (EEG) with high-temporal-resolution can also provide valuable evidence supporting these two distinct components. Firstly, theta activity was suggested to play a central role in learning and behavioral adjustments ^24,25^. Recent study has shown that midfrontal theta increased when individuals reassessed their judgments after observing others make inconsistent decisions ^26^. Of note, the study found that midfrontal theta activity increased as the expected reward grew ^27^, suggesting that theta activity was involved in processing reward amounts. Thus, we hypothesize that social influence would alter late theta activity related to the weight of amount, with its corresponding time window occurring after option onset. Secondly, study found that the starting point reflecting early motor preparation in decision choice ^28^. Moreover, central beta activity was associated with motor preparation ^29^ and could also predicted the decision starting point ^30^. Therefore, we hypothesize that social influence would change early beta activity related to the starting point, with the corresponding time window occurring before option onset. In summary, as illustrated in Figure 1, we would combine DDM and EEG, employing multivariate pattern analysis to provide convergent evidence to support the two-components hypothesis.

**Figure 1.**
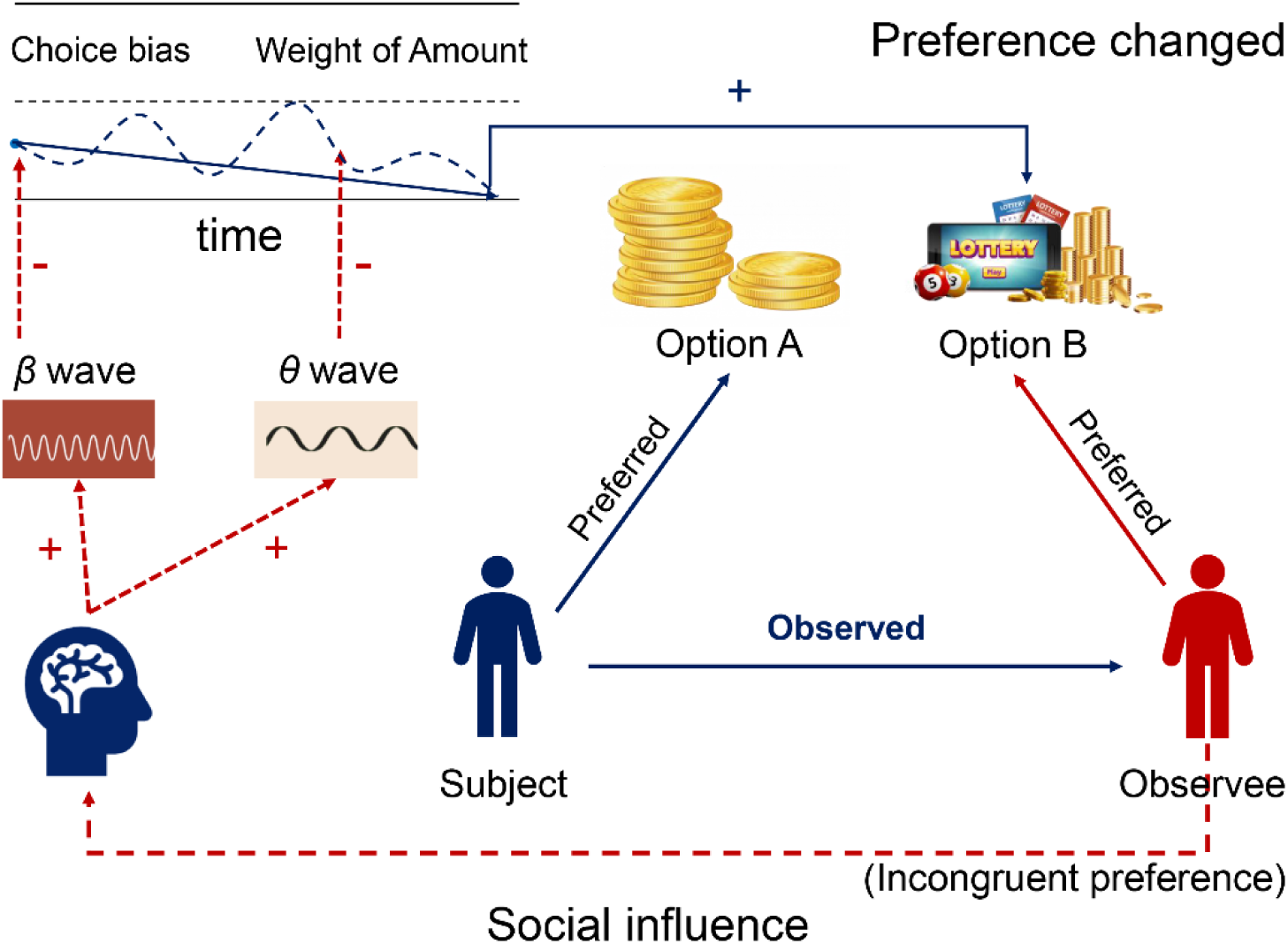
Overview of the theoretical framework of this study. After individual observes other with risk preferences inconsistent with their own, he exhibits changes in *β* activity, accompanied by a shift in choice bias toward the option more frequently chosen post-observation. Then *θ* activity changes occur, leading to a shift in the weighting of monetary amounts, which biases subjective value toward the same option. Ultimately, this results in a change in the individual’s risk preference.

## 2 Results

### 2.1 Risk decision-making performance and HDDM fitting results

First, to know the risk preference distribution of the subjects, we summarized each participant’s risk preference prior to observation. Consistent with prior studies ^5,31^, the majority were found to be risk-averse (see Figure 3.A). Subsequently, to investigate whether a social influence effect existed, a mixed-design repeated-measures ANOVA was conducted on the risk preference consistency (Stage: pre-observation vs. post-observation × Group: Incon vs. Con). The results revealed that the main effect of stage was significant (*F*(1,57) = 8.05, *p* = 0.006, *η²p* = 0.12), as was the main effect of group (*F*(1,57) = 11.69, *p* = 0.001, *η²p* = 0.17). Furthermore, there was a significant interaction between stage and group (*F*(1,57) = 14.55, *p* < 0.001, *η²p* = 0.20). Simple effects analysis revealed that (see Figure 3.B), the main effect of group was significant in the post-observation stage (*F*(1,57) = 22.71, *p* < 0.001) but not in the pre-observation stage (*F*(1,57) = 1.86, *p* = 0.175). Specifically, in the post-observation phase, the Incon group exhibited more negative risk preference consistency (*M* = -0.02, *SE* = 0.03) compared to the Con group (*M* = 0.12, *SE* = 0.02).

**Figure 2.**
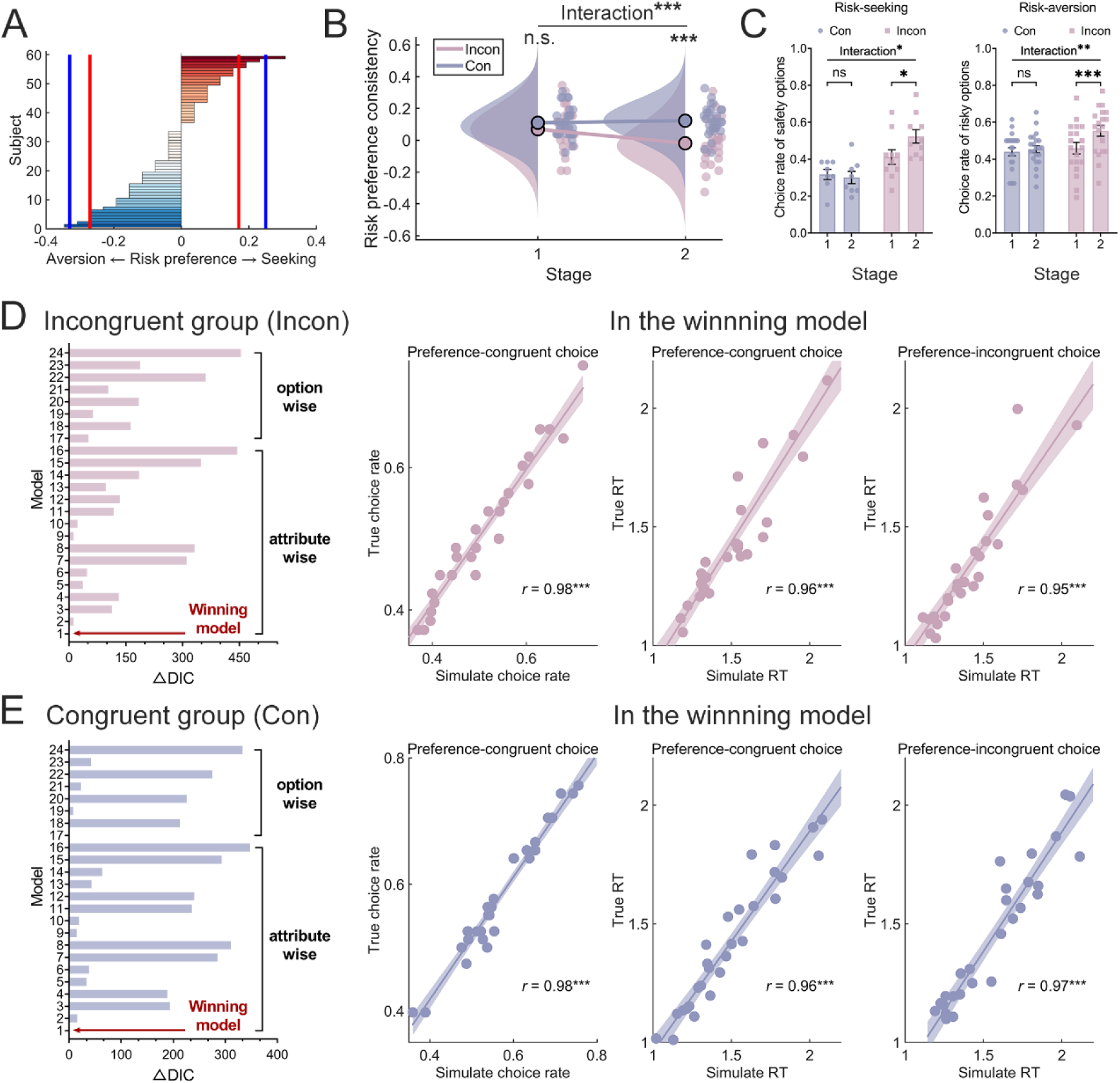
Risk preference and HDDM fitting results. **(A)** The distribution of risk preferences for all observers (the solid lines in the figure represented the degree of risk preference of the four observes. blue: male; red: female). **(B)** The social influence effect. Stage 1 and 2 referred to pre-observation and post-observation, respectively. **(C)** The change in choice rates for individuals with different risk preferences due to social influence. **(D)** The difference between the smallest DIC and DIC of other competitive models in the Incon group. And the correlations between the simulated and observed data in the Incon group in regrade to the choice rate of preference-congruent options (left), the RT of preference-congruent choices(middle), and the RT of preference-incongruent choices (right). **(E)** The difference between the smallest DIC and DIC of other competitive models in the Con group. And the correlations between the simulated and observed data in the Con group in regrade to the choice rate of preference-congruent options (left), the RT of preference-congruent choices (middle), and the RT of preference-incongruent choices (right) Each dot represented one participant. Preference-incongruent choices referred to the options increased after observation. Shaded areas were the 95% confidence intervals (*: *p* < 0.05, **: *p* < 0.01, ***: *p* < 0.001, the same as below).

**Figure 3.**
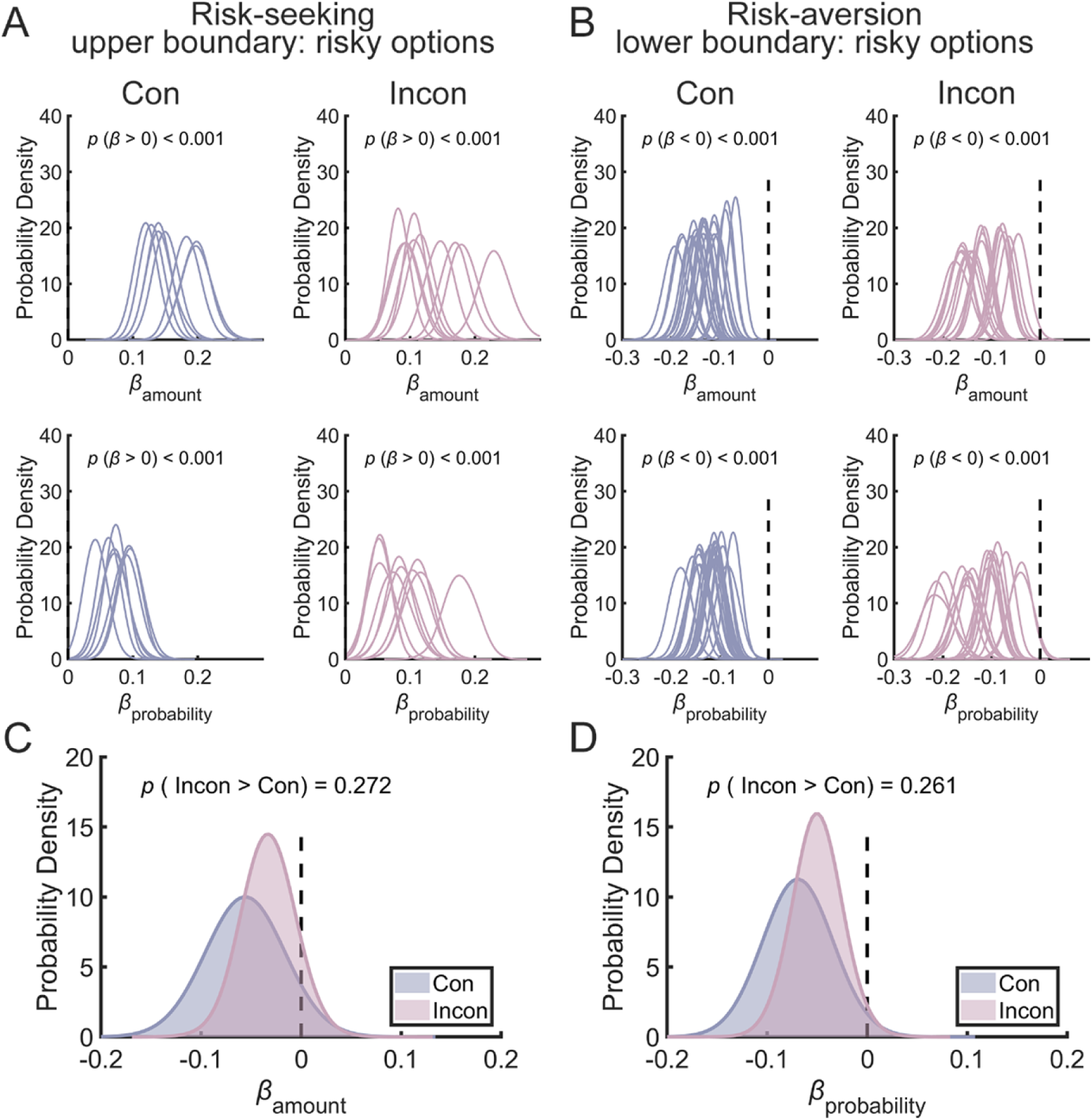
Amount and probability drove value accumulation in risky decision-making. (A) Posterior probability distributions of amount and probability contributions to value accumulation for risk-seeking participant. (B) Posterior probability distributions of amount and probability contributions to value accumulation for risk-aversion participant. (C) Comparison of the distribution of amount contributions to value accumulation between the Incon and Con groups. (D) Comparison of the distribution of probability contributions to value accumulation between the Incon and Con groups.

To elucidate how social influence altered the choice rates for different options, a repeated-measures ANOVA (Stage: pre-observation vs. post-observation × Group: Incon vs. Con) was conducted simultaneously on the choice rate of the safe option among risk-seeking individuals and on the choice rate of the risky option among risk-aversion individuals. For risk-seeking individuals, a significant interaction effect between stage and group was observed (*F*(1,16) = 6.71, *p* = 0.02, *η²p* = 0.30). Simple effects analysis revealed that (see Figure 3.C left), the main effect of group was significant in the post-observation stage (*F*(1,16) = 19.39, *p* < 0.001) but not in the pre-observation stage (*F*(1,16) = 3.45, *p* = 0.082). Similarly, for risk-aversion individuals, a significant interaction effect between stage and group was observed (*F*(1,39) = 7.58, *p* = 0.009, *η²p* = 0.16). Simple effects analysis revealed that (see Figure 3.C right), the main effect of group was significant in the post-observation stage (*F*(1,39) = 8.53, *p* = 0.006) but not in the pre-observation stage (*F*(1,39) = 0.26, *p* = 0.614). Overall, social influence led individuals with different risk preferences to choose more options incongruent their original preferences after observation.

Based on the aforementioned results, the lower boundary in the HDDM was defined as the option that was more frequently chosen by participants in the Incon group after observation (corresponding to the safe option for risk-seeking individuals and the risky option for risk-aversion individuals). In both the Con and Incon groups, we constructed 24 HDDM models to investigate if risk decision-making could be captured by the decision parameters of drift rate (*v*), starting point (*z*), and threshold (*a*). As depicted in Figure 3.D, we found that model #1 had the smallest DIC in Incon group, with the difference from the second smallest DIC exceeding 10 (Incon: ΔDIC = 12.05). Given that previous research suggested that a difference greater than 10 is considered significant, model #1 should be considered the winning model. Next, in Model 1, we simulated participants’ choice rates and RT for various decisions. The simulated values showed a high correlation with the actual data, with all correlation coefficients (*r*) exceeding 0.95, indicating that Model 1 fit the observed data very well. Similarly, we found that model #1 had the smallest DIC in Con group (see Figure 3.E). Although the difference from the second smallest DIC did not exceed 10 (Con: ΔDIC = -1.07), the simulated choice rates and RT for various decisions showed a high correlation with the actual data, with all correlation coefficients (*r*) exceeding 0.96, indicating that Model 1 also fit the observed data very well.

### 2.2 The amount and probability drive value accumulation in risk decision-making

To verify that both the monetary amount and probability had main effects on the accumulation of value in risk decision-making for each trial, we first examined the distribution of these two variables separately among risk-seeking and risk-aversion individuals. For risk-seeking individuals, since the risky option was defined as the upper boundary in the HDDM, the analysis revealed that in both groups, the probability of the contribution values of amount and probability to value accumulation exceeding zero was greater than 99% (see Figure 4.A). For risk-aversion individuals, since the risky option was defined as the lower boundary in the HDDM, the probability of the contribution values of amount and probability to value accumulation being less than zero exceeded 99% in both groups (see Figure 4.B). These results aligned with common sense: as the amount or probability increased, individuals were more likely to choose the risky option and thus confirmed that both monetary amount and probability exhibited significant main effects on the accumulation of value during risk decision-making. Finally, we found that the probability that the contribution of amount to value accumulation was greater in the incongruent group (mean highest density interval (HDI) = -0.033, 95%HDI = [-0.084, 0.020]) than in the congruent group (mean HDI = -0.056, 95%HDI = [-0.109, -0.004]) was 27%, which was not statistically significant. Similarly, the probability for probability contribution was 26% (Incon: mean HDI = -0.050, 95%HDI = [-0.097, -0.028]; Con: mean HDI = -0.069, 95%HDI = [-0.031, -0.107]), also not significant.

**Figure 4.**
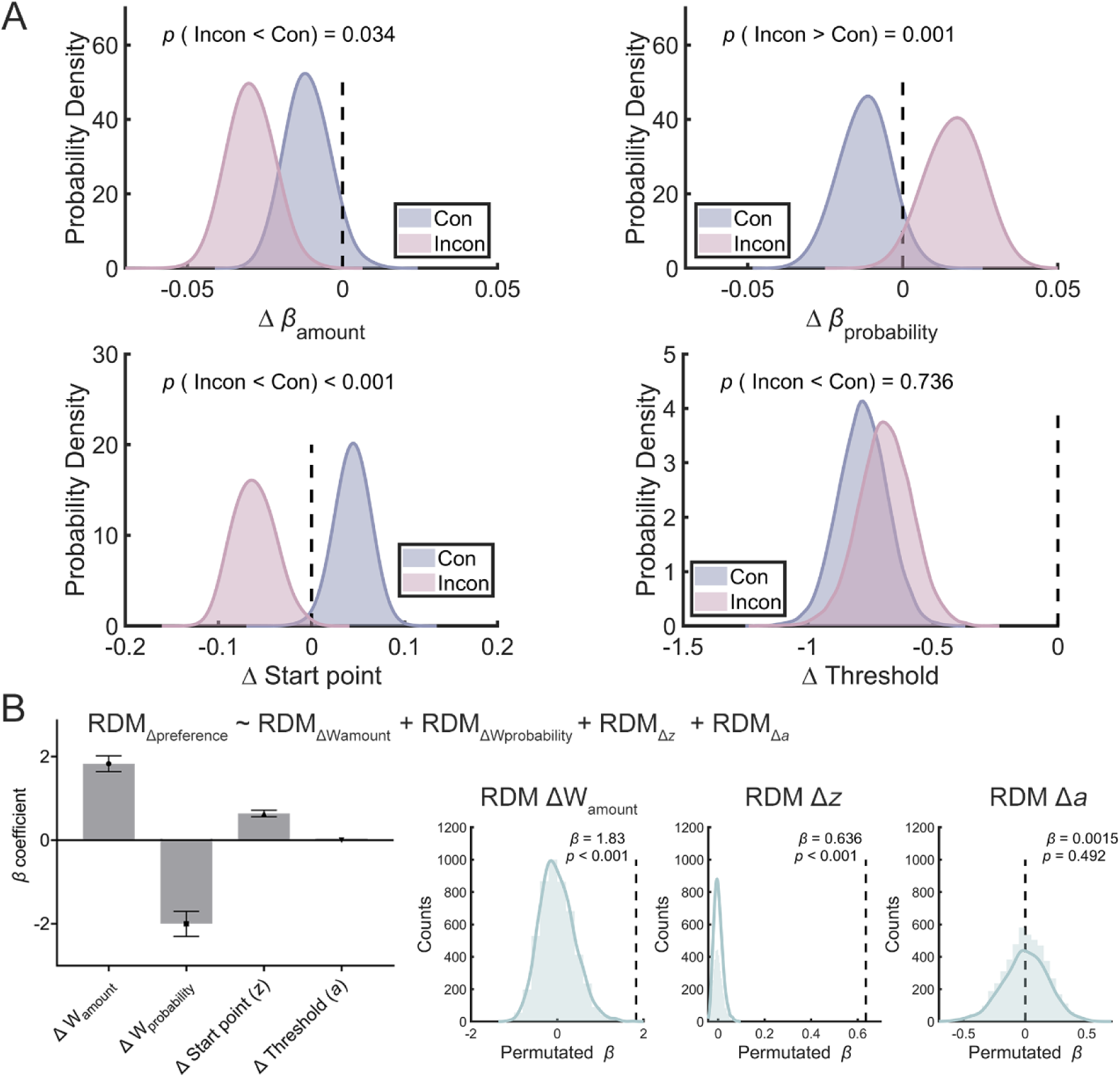
The key processes modulated by social influence. (A) Between-group comparison of changes in different parameters (post-observation minus pre-observation). (B) Inter-subject representational similarity analysis revealed the associations between changes in each parameter and changes in risk preference consistency, along with permutation test results for all positive prediction coefficients.

### 2.3 Social influence modulated weight of amount in value accumulation and starting point

To investigate how social influence altered the risk decision-making process, we compared the changes (post-observation minus pre-observation) in weight of amount (W_amount_), weight of probability (ΔW_probability_), decision starting point (*z*), and decision threshold (*a*) between groups (see Figure 5.A). In the Incon group, compared to pre-observation (Stage 1), the weight of the amount was significantly more negative (the probability of ΔW_amount_ being negative > 99%). However, in the Con group, this was not the case (the probability of ΔW_amount_ being negative = 94.54%). Crucially, ΔW_amount_ in the Incon group was significantly more negative than in the Con group (posterior probabilities = 96.6%). Also, the weight of probability was strengthened after observation in both the Incon group (the probability of ΔW_probability_ being positive > 98%) but was weaken in the Con group (the probability of ΔW_probability_ being negative > 96%). The difference between the two groups was also significant (posterior probabilities > 99%).

**Figure 5.**
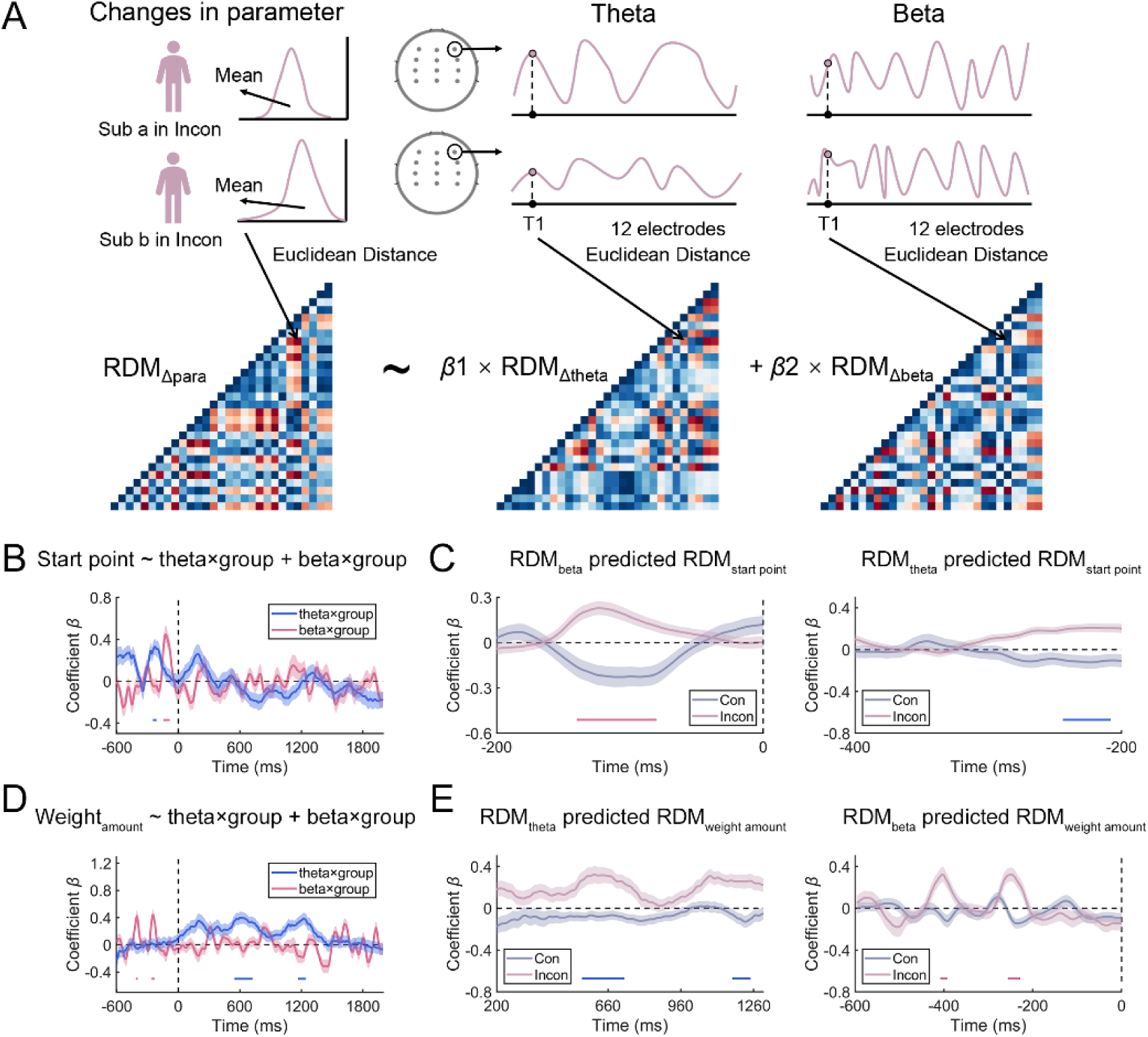
Inter-subjects similarity analysis for the neural activities. (A) Schematic illustration of inter-subject RSA analysis at the neural level. (B) The prediction of interaction between RDM_theta_ and RDM_beta_ on RDM_starting point_ at each sampling point. (C) Prediction of the RDM_starting point_ by RDM_theta_ and RDM_beta_ in each group at sampling points showing significant interaction effects (during -144 to -80 ms). (D) The prediction of interaction between RDM_theta_ and RDM_beta_ on RDM_weight amount_ at each sampling point. (E) Prediction of the RDM_weight amount_ by RDM_theta_ and RDM_beta_ in each group at sampling points showing significant interaction effects (during 548 to 728 ms and 1168 to 1248 ms).

Similarly, relative to pre-observation, the starting point significantly decreased in the Incon group (posterior probabilities > 99%) but increased in the Con group (posterior probabilities > 99%). The relative shift in the starting point between the two groups was significant (posterior probabilities > 99%). The decision threshold significantly decreased across both groups (posterior probabilities > 99%), yet there was no significant difference between the two groups (posterior probabilities = 73.56%). The mean HDI and 95% HDI for each parameter were presented in Supplementary Table S2, and the distributions of parameter changes for each participant were shown in Supplementary Figure S1.

Finally, to further reveal the decision-making processes associated with behavioral changes, we employed inter-subject representational similarity analysis (IS-RSA) in all participants. Euclidean distances were used to construct representational dissimilarity matrices (RDMs) for the above parameters (the mean value of each distribution), which were then used to predict behavioral RDMs via a general linear model (GLM). The results showed that the beta coefficient of ΔW_amount_ was significantly greater than 0 (*β* = 1.83, *SE* = 0.19, *p* < 0.001), the beta coefficient of ΔW_probability_ was negative (*β* = -2.00, *SE* = 0.30), the beta coefficient of Δ_starting point_ was significantly greater than 0 (*β* = 0.64, *SE* = 0.08, *p* < 0.001) but the beta coefficient of Δ_threshold_ was positive but not significantly greater than 0 (*β* = 0.002, *SE* = 0.007, *p* = 0.492) (see Figure 5.B). These results indicated that social influence primarily changed the risk decision-making process by modulating the weight of amount and the decision starting point.

### 2.4 Dynamic neural activities underlying the changed DDM parameters

To identify neural activities associated with the two key parameters, we further applied IS-RSA to examine how neural patterns dynamically represented HDDM parameters over time. As shown in Figure 6A, we constructed parameter RDMs separately for the Incon and Con groups. By calculating the Euclidean distances across 12 electrodes between different participants, we generated theta and beta RDMs and then established GLMs. First, we used the GLM (dummy code: Con group = 1; Incon group = 2) to simultaneously assess the interactions of RDM_theta_ × group and RDM_beta_ × group on predicting RDM_starting point_. The results revealed group differences of theta and beta activity before option presentation in predicting the starting point, with beta showing a greater number of significant sampling points between different groups (theta × group: 10 consecutive sampling points, corresponding time window: -248 to -208 ms, *β_min_* = 0.29, *t*_min_ = 4.00, *p*_FWE max_ = 0.045; beta × group: 16 consecutive sampling points, corresponding time window: -144 to -80 ms, *β_min_*= 0.32, *t*_min_ = 4.08, *p*_FWE max_ = 0.032, see Figure 6.B). According to the prior hypothesis, we further examined how RDM_beta_ within the −144 to -80 ms window predicted the RDM_starting point_ in each group separately. Simple effects analysis revealed that, after controlling for RDM_theta_, RDM_beta_ positively predicted the RDM_starting point_ in the Incon group (*β_min_*= 0.11, *t*_min_ = 2.84, *p*_FWE max_ = 0.075), whereas the prediction was negative in the Con group (*β_max_* = -0.16, *t*_max_ = -2.47, *p*_FWE min_ = 0.008, see Figure 6.C).

**Figure 6.**
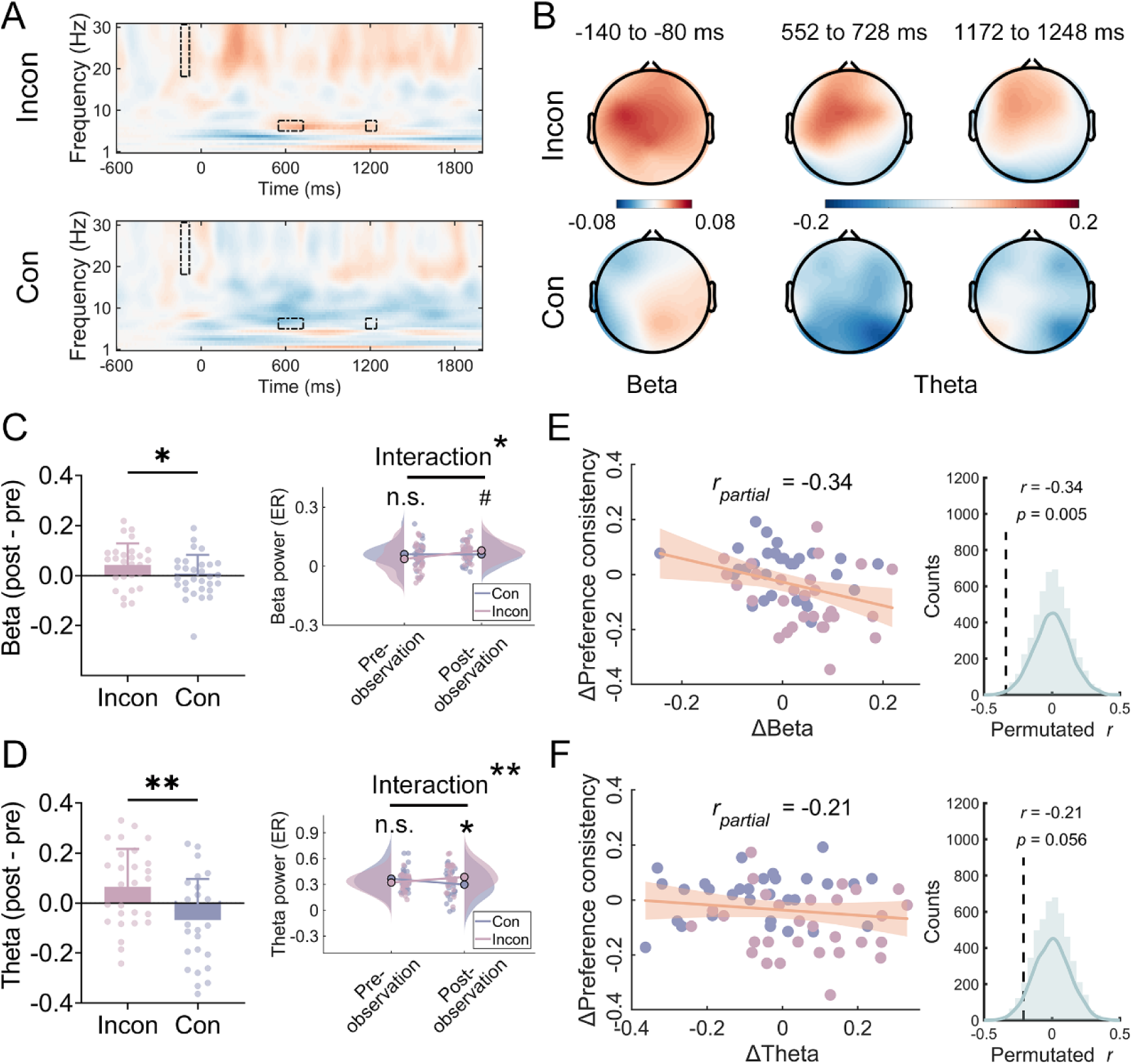
Univariate analysis for the neural activities. **(A)** The mean time-frequency representations in 1–30 Hz across selected electrodes. **(B)** Topographic maps of averaged neural data (post-observation minus pre-observation) for beta power (during -144 to -80 ms) and theta power (during 548 to 728 ms and 1168 to 1248 ms) in each group. **(C)** Group comparison of social influence–induced change in beta power. **(D)** Group comparison of social influence–induced change in theta power. **(E)** Correlation between beta and behavior. **(F)** Correlation between theta and behavior. #: 0.05 < *p* < 0.1.

Then, we used the GLM to simultaneously assess the interactions of RDM_theta_ × group and RDM_beta_ × group on predicting RDM_weight amount_. The results revealed group differences of theta and beta activity after option presentation in predicting the weight amount, with theta showing a greater number of significant sampling points between different groups (theta × group: 45 and 20 consecutive sampling points in two periods, corresponding time window: 548 to 728 ms and 1168 to 1248 ms, *β_min_* = 0.32, *t*_min_ = 3.98, *p*_FWE max_ = 0.049; beta × group: 5 and 8 consecutive sampling points in two periods, corresponding time window: -412 to -392 ms and -260 to -228 ms, *β_min_* = 0.35, *t*_min_ = 4.02, *p*_FWE max_ = 0.041, see Figure 6.D). According to the prior hypothesis, we further examined how RDM_theta_ within the 548 to 728 ms and 1168 to 1248 ms window predicted the RDM_weight amount_ in each group separately. Simple effects analysis revealed that, after controlling for RDM_beta_, RDM_theta_ positively predicted the RDM_weight amount_ in the Incon group (*β_min_* = 0.24 *t*_min_ = 3.35, *p*_FWE max_ = 0.057), whereas the prediction was negative in the Con group (*β_max_* = -0.16, *t*_max_ = -1.14, *p*_FWE min_ = 1.65, see Figure 6.B).

### 2.5 Social influence shaped the theta and beta activity

Figure 7. A displayed the mean difference (post-observation minus pre-observation) across the selected electrodes (12 in total) at 1–30 Hz time-frequency representations in the Incon group (top) and Con group (bottom). Topographical maps revealed an increasing trend in both beta and theta power in the Incon group under social influence (see Figure 7.B). To characterize this trend, we conducted independent samples T-test to compare the changes in beta and theta power between groups. Results showed that the increase in beta power in the Incon group was marginally significantly greater than in the Con group (*t*(57) = 1.99, *p* = 0.052, see Figure 7.C (left); similarly, the increase in theta power was significantly greater in the Incon group (*t*(57) = 3.19, *p* = 0.002, see Figure 7.D (left)). To more comprehensively account for the non-independence of repeated measures data, we further employed the linear mixed-effects model where participant was incorporated as a random intercept to examine the interaction between group and time (dummy code: Con group = 1; Incon group = 2) and time (dummy code: pre-observation = 1; post-observation = 2). For beta (see Figure 7.C (right)), we found a significant interaction between group and time (*β* = 0.04, *t*(114) = 2.18, *p* = 0.03). Simple effects analysis revealed that before observation, there was no significant difference between groups (Incon: *M* = 0.04, *SE* = 0.01; Con: *M* = 0.06, *SE* = 0.01; *β* = -0.02, *t*(57) = -1.48, *p* = 0.15), whereas after observation, the beta power in the Incon group was marginally significantly higher than the Con group (Incon: *M* = 0.08, *SE* = 0.01; Con: *M* = 0.06, *SE* = 0.01; *β* = 0.02, *t*(57) = 1.68, *p* = 0.099). For theta (see Figure 7.D (right)), we found a significant interaction between group and time (*β* = 0.13, *t*(114) = 3.25, *p* = 0.002). Simple effects analysis revealed that before observation, there was no significant difference between groups (Incon: *M* = 0.32, *SE* = 0.02; Con: *M* = 0.37, *SE* = 0.02; *β* = -0.04, *t*(57) = -1.57, *p* = 0.12), whereas after observation, the theta power in the Incon group was significantly higher than the Con group (Incon: *M* = 0.39, *SE* = 0.03; Con: *M* = 0.30, *SE* = 0.03; *β* = 0.09, *t*(57) = 2.31, *p* = 0.02).

**Figure 7.**
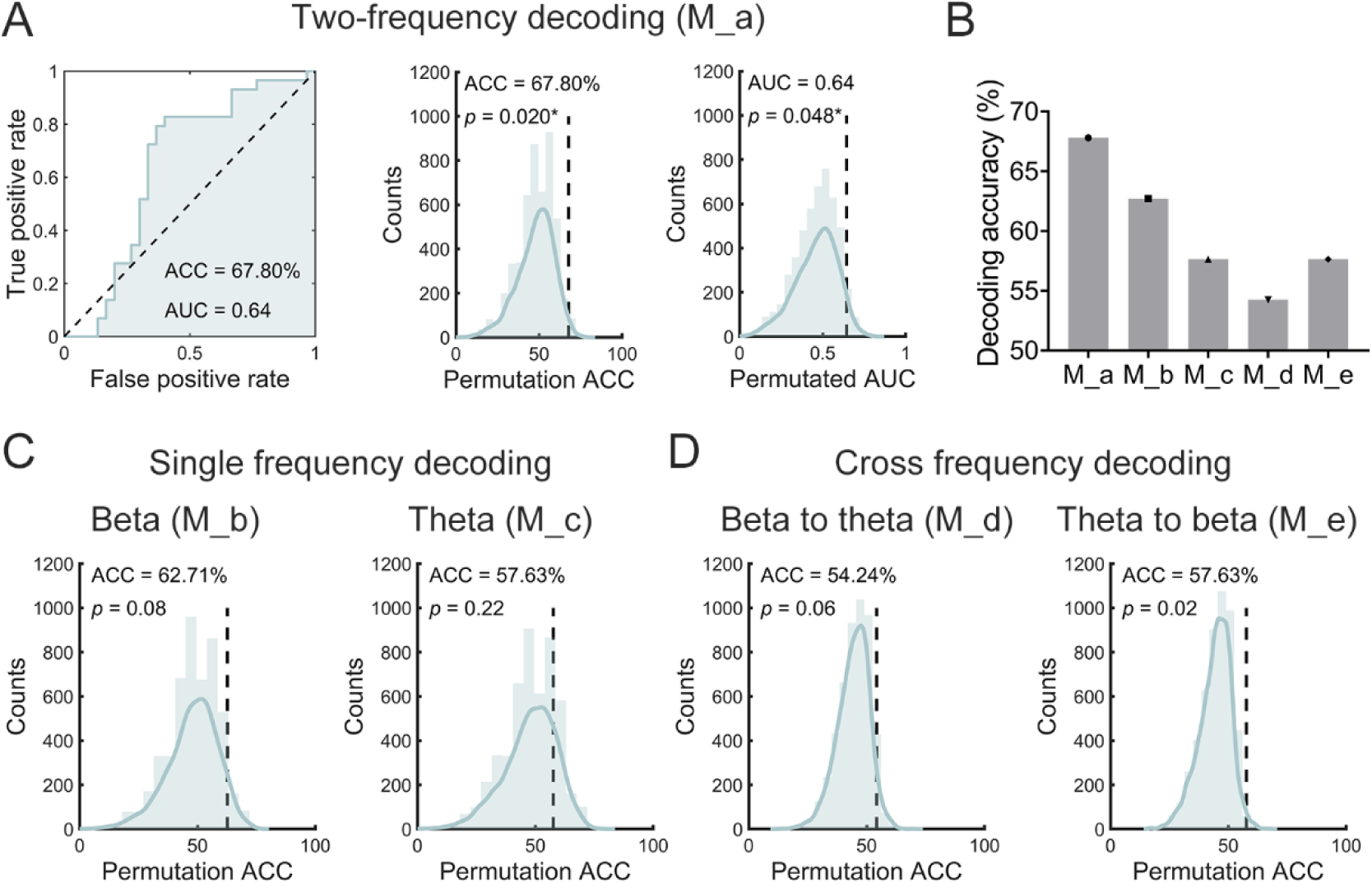
Multivariate analysis for the neural activities. (A) Performance of classifying groups when theta and beta activities were used jointly as features. (B) Classification performance of different models. (C) Permutation test results for classification accuracy using single neural activities. (D) Permutation test results for cross-frequency neural activity classification accuracy.

After pooling all participants and controlling for initial risk preference, partial correlation analyses showed that changes in beta activity were negatively correlated with changes in risk preference consistency (*r* = -0.34, *p* = 0.005), while changes in theta activity were marginally negatively correlated with changes in risk preference consistency (*r* = -0.21, *p* = 0.056).

### 2.6 Theta and beta in the different stage supported two distinct components

To further verify that theta and beta supported different stages yet jointly contributed to decision-making rather than simply covarying, we employed a machine learning approach to examine their classification performance for distinguishing groups when combined, when used separately, and in cross-frequency prediction. First, in the combined model (M_a), we averaged the beta power across all sampling points significantly predicting changes in the starting point, and the theta power that significantly predicting changes in weight of amount, resulting in a 59 × 12 feature matrix. This matrix was then input into an SVM classifier and the classification accuracy for distinguishing the two groups was 67.8%, with the area under the ROC curve (AUC) reaching 0.64. The permutation test with 5000 iterations revealed that both the classification accuracy and the AUC were statistically significant, with *p*-values less than 0.05 (see Figure 8.A). In the single models, we averaged only the beta (M_b) or theta (M_c) at significant sampling points to form 59×12 feature matrices and assessed their classification performance for group. In the cross-frequency models, we first constructed the same 59×12 feature matrices, then trained the model with beta features and tested classification using theta features (M_d), and vice versa (M_e). Among all classification results, the combined model (M_a) yielded the best performance (see Figure 8.B). Notably, in the single models, although classification accuracy exceeded 60%, it was not significant (see Figure 8.C); in the cross-frequency models, accuracy was marginally or significantly above chance but remained below 60% (see Figure 8.D). Overall, only when theta and beta activities were considered together did the model perform most comprehensively, supporting that these two neural activities not only occur at different stages but also support distinct functions.

**Figure 8.**
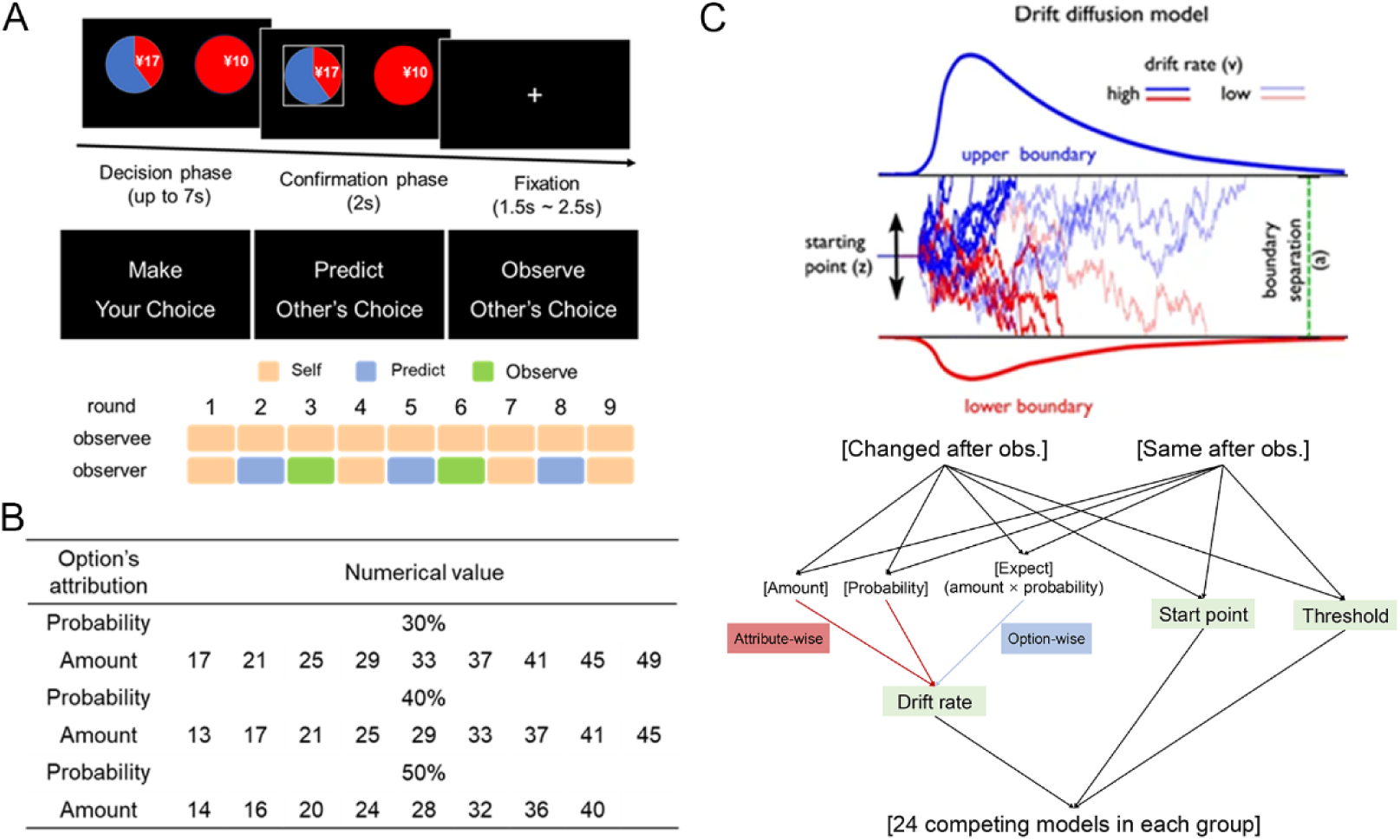
The details of experiment. (A) The procedure of trial (top), the different instructions in experiment (middle) and overall schedule for the observe and observer (bottom). (B) The specific combinations of amount and probability in the experiment and the amounts were manipulated by increase by 1, decrease by 1, or remain unchanged. (C) Decision-making process assumed by the drift diffusion model (top, from: Pedersen et al., 2017, *Psychonomic bulletin & review*). All hypothesized model in the present study (bottom). First, six possibilities were defined based on whether the weights of amount, probability, or expected value (amount × probability) in the drift rate changed before and after observation. Then, four additional possibilities were defined based on whether the starting point and decision threshold changed before and after observation. Therefore, 24 competing models were defined for each group.

## 3 Discussions

This study grouped participants were into consistent (Con) and inconsistent (Incon) groups based on whether their initial risk preferences aligned with those of the observes and use the DDM and time-frequency analysis to examine the potential components altered by social influence during risk decision-making. After observation, the Incon group showed decreased risk preference consistency: risk-seeking individuals in chose more safety options, while risk-averse individuals chose more risky options, compared to the Con group (to facilitate subsequent discussion, we will uniformly refer to these options as preference-inconsistent choices). This replicated findings of previous study ^5^.

In the DDM framework (lower boundary: options increased after observation in Incon group), we found that attribute-wise model provided a better fit than option-wise model in capturing how social influence altered individual risk decision-making. This was the first to reveal that social influence altered individuals’ risk decision-making in the attribute-wise process instead of option-wise. The results is consistent with recent study providing supporting evidence that these attributes are compared independently in risky decision-making ^12^. It was suggested that amount was considered before probability in risk decision-making ^14^. A recent study has showed that attribute-wise models fit intertemporal decision-making better than option-wise models ^23^. We propose that an attribute-wise heuristic may allow participants to identify their preferred options effortlessly under time constraints.

Further, according to the best-fitting attribute-wise model, we discovered that social influence led to a negative increase in the weight of the amount. Consistent with prior research, these findings demonstrated how observing others’ choices could modulate value computation processes ^18,32–34^. In general, others’ goals could influence one’s goals, preferences, and choices ^8,9^, prioritizing attributes consistent with these new goals once changes occurred ^11^. For example, in dietary decision-making, focusing on health goals led to healthier choices by increasing the weight of healthy attributes in value computations ^35^. Since probability might often be completely disregarded when decisions were made under uncertainty ^13^ and priority heuristic theory claimed that prioritization of amount was over probability ^14^, our results implied that the amount served as a critical cue for participants to infer and learn others’ risk decision-making behaviors. This helps explaining why, in the second stage, only the weight of amount showed both between-group differences and positively representing the changes in risk preference consistency—but not observed for probability.

Additionally, social influence caused smaller starting point and negatively increased weight of amount in value accumulation in the Incon group, revealing social influence changed two decision stages. First, observing others’ decisions reduced the starting point (*z*) of decision-making in the incongruent group. The starting point reflected the initial choice bias toward one option, which existed before processing each attribute during risk decision-making ^20^. It was demonstrated that choice bias could be shaped. A recent study using the DDM model found that observing an antisocial peer’s moral decision made the starting point more biased toward the harmful option ^18^. Since the lower boundary represented the preference-inconsistent choice in the study, the smaller starting point change suggested that social influence caused the incongruent group to favor preference-inconsistent choices before processing any attribute. In fact, individuals often imitate the behavior of others outside of conscious awareness and intent ^16,17^, driving an individual to activate an updated state by imitating the observee (i.e., making a similar choice). Also, individuals exhibited a status quo bias, referring to a tendency to maintain the current state of affairs ^36^. Namely, individuals would like to maintain learned state in the new situation (i.e., repeating the similar choice). These option-independent states may collectively drive the choice bias first changed by social influence.

At neural level, inter-subject RSA analysis revealed that in the Incon group, changes in later theta activity (500 ms after option onset) more positively represented changes in the weight of the amount in value accumulation, while changes in early beta activity (prior to option onset) more positively represented changes in starting point. More specifically, linear mixed-effects modeling revealed that social influence led to significantly greater post-observation theta and beta activity in the Incon group compared to the Con group. Importantly, the classification performed best only when theta and beta were combined as features. These results supported that theta and beta reflected two distinct components, further revealing how social influence modulates the decision-making process.

We found that in the Incon group, changes in later theta activity (500 ms after option onset) more positively represented changes in the weight of the amount attribute during value accumulation. What’s more, linear mixed-effects modeling revealed that social influence led to significantly greater post-observation theta activity in the Incon group compared to the Con group. Previous studies showed that observing decisions inconsistent with personal preferences increased theta activity when individuals re-evaluated faces ^26^ or brands ^37^. In general, inconsistencies provoked aversive arousal. This led individuals to control their behavior intentionally to reduce discomfort ^38^. Several studies in non-social contexts had demonstrated that increased theta activity was linked to enhanced cognitive control and conflict resolution ^24,39^. Thus, the increased theta in the Incon group suggested that social influence had individuals to exert more effort to alleviate the discomfort caused by inconsistency. Of note, previous study found that theta activity increased with expected rewards ^27^, demonstrating its involvement in processing amounts. This helped explaining why changes in theta activity represented changes in the weight of amounts during value accumulation.

Besides, changes in early beta activity (prior to option onset) more positively represented changes in starting point and the Incon group showed greater beta activity than the Cong group after observation. Previous studies found that observing others’ inconsistent decisions caused decreased beta power. They suggested that reduced beta activity reflected negative feedback when others’ decisions conflicted with one’s own ^37,40^. However, beta activity in our study likely served a different function especially considering that the beta activity in our study emerged before the option onset (outside the baseline period) but after the option onset in previous study. Recent research has showed that motor-related beta activity predicted the starting point ^30^, which aligns with our results. Other studies found that, the starting point influenced decision choice through early motor preparation ^28^, suggesting that beta activity in our study reflected motor control ^29,41,42^. Previous studies also showed that inconsistencies with the correct response increased beta power in the next trial, indicating inhibited motor processes ^43,44^. The increased beta power in the Incon group suggested that social influence suppressed motor processes for preference-consistent choices during early decision-making.

Since only when theta and beta activities were considered together in machine learning the did the classification model perform best, the neural results further convergently supported the hypothesis that social influence changed risk decision-making through two distinct and sequent stages. However, this study had several limitations. First, all participants were from China, where cultural emphasis on collectivism was higher compared to Western cultures. Previous research showed that the effect size of social influence on buying decisions was positively associated with collectivism ^45^. Therefore, the findings might not have been generalizable across cultures. Second, the study did not account for the potential influence of personality traits on the results. While agreeableness did not affect the behavioral impact of social influence, it was found to modulate neural processing during the observation of others’ decisions ^40^. This suggested caution when generalizing the neural findings of this study. Finally, to ensure uniformity in the observed decisions, observees’ choices were pre-recorded and later presented to observers. This approach limited the revealed neural activities to a single-brain perspective. In contrast, real-time social interactions during observational learning had been shown to increase alpha-mu synchronization between observers and observees ^46^. Given that this study only focused on the results of social influence on risk decision-making, future research could employ real-time hyper-scanning techniques to further focus on the observation and prediction phases, investigating the processes of social learning in the social influence for a more comprehensive understanding.

In summary, via computational modeling and multivariate analysis of EEG, we explored how social influence modulated the decision-making process. Using the DDM, we first discovered that social influence acted via an attribute-wise way. We further established that this process unfolded in two distinct components: the prioritization of amount in value accumulation and choice bias. The two components were further supported by neural evidence. As changes in later theta activity (500 ms after option onset) more positively represented changes in the weight of the amount attribute during value accumulation, while changes in early beta activity (prior to option onset) more positively represented changes in starting point. In addition to the changes of theta and beta could collectively distinguish different groups, enhanced theta suggested increased efforts to adapt to inconsistency and enhanced beta suggested motor suppression during the initiation of decision-making. All the results convergently supported our theoretical hypothesis regarding the dynamics of social influence (see Figure 1), helping understanding the behavioral contagion during learning about another agent’s risk-preferences acts. It also offers implications for future economic research. For instance, social influence in risk decision-making can contribute to the formation and collapse of financial bubbles. However, the specific roles and interplay of the two mechanisms in this phenomenon warrant further investigation.

## 4 Methods

### 4.1 Participants

G*Power v. 3.1 software ^47^ suggested that a minimum of 54 participants would be needed for a 2 x 2 mixed-design repeated-measures ANOVA, including an interaction, to detect a moderate effect size (*f* = 0.25) with default alpha and power values (*α* = 0.05 and 1-*β* = 0.95, respectively). Consequently, the study recruited 64 participants (mean age 22.9 ± 2.7 years), including 4 observees (2 males) and 60 observers (24 males). Observers were paired with observees of the same gender based on their self-reported risk preferences. Pairs with same risk preferences were categorized as the risk preference consistent group (Con group), where risk-seeking observees were matched with risk-seeking observers and risk-averse observees with risk-averse observers. Conversely, pairs with opposite preferences formed the risk preference inconsistent group (Incon group), with risk-seeking observees matched with risk-averse observers and vice versa. Ultimately, there were 30 participants in the Con group and 29 in the Incon group (one excluded due to a preference change greater than 3SD from the mean). There were no significant differences in gender (*χ*² = 0.91, *p* = 0.34) or age (*t*(57) = 0.76, *p* = 0.45). All participants were right-handed, had normal or corrected-to-normal vision, no history of psychiatric disorders, and had signed an informed consent form. They received a stipend upon completion. The study was approved by the Human Research Protection Committee of East China Normal University (No. HR 429-2020).

### 4.2 Experimental procedure and task

Initially, the researcher assigned roles to the participants as either observees or observers. Immediately following this, instructions were provided, and an EEG cap was fitted on each participant. Before the main experiment, participants engaged in a 4-trial practice session to ensure they fully understood and were comfortable with the task.

This task was adapted from a paradigm used in the previous study ^5^. As illustrated in Figure 2.A (top), each trial comprised two phases: a decision phase followed by a confirmation phase. In the decision phase, two pie charts appeared on the screen simultaneously, representing a safe option and a risky option. The red portion of each pie chart indicated the probability of winning money, while the figures within the charts displayed the monetary amounts associated with each option. The positions of the pie charts were randomized on the screen. Participants made their choices by pressing the “z” key for the left option and the “c” key for the right option, with up to 7 seconds allowed for making a decision.

If no selection was made or after a choice was made, the confirmation phase began. If the risky option was selected, participants would gamble the amount shown for that risk in the chart; if the safe option was selected, they would receive ¥10 directly. During this phase, a white box highlighted the chosen pie chart for 2 seconds to confirm the selection. The trial concluded with a fixation point displayed for a random duration between 1.5 and 2.5 seconds.

Participants were required to complete nine rounds of the task, with each round consisting of 28 trials, including two probe trials. In these probe trials, both options were safe but offered different monetary amounts. These trials served to assess participant engagement and were not included in the final analysis. In the remaining 26 trials of each round, both the probability and the amount of money varied (see Figure 2.B). To enhance the realism of the scenarios, random noise was added to the amounts in each trial, which could increase by 1, decrease by 1, or remain unchanged.

As depicted in Figure 2.A (middle), observers not only made their own choices but also had tasks involving prediction and observation of another’s choices. Under the prediction instruction, observers needed to guess the decisions that the observee would make during the decision phase. During the confirmation phase, only the option predicted by the observer was displayed, without revealing whether the prediction was accurate. Under the observation instruction, observers were required to carefully watch the choices made by the observee during specific rounds (third and sixth rounds of the observee’s task). This was the only condition under which observers could learn about the observee’s risk preferences. Figure 2.A (bottom) illustrated the sequence of different instructions given to both the observee and the observer.

### 4.3 Data collection

#### 4.3.1 Behavioral data

Presentation of task stimuli, control of task flow, and collection of behavioral data were performed by PsychoPy v3.0 ^48^. The library is based on the python platform.

#### 4.3.2 EEG data

A 32-channel EEG cap was used to record EEG data in this study (Compumedics NeuroScan). The electrodes on this EEG cap were distributed according to the International 10/20 system. Ocular electrical signals were recorded by four additional electrodes. Two of the horizontal electrodes were placed on either side of the outer corners of the eyes in the left and right eyes, while the other two vertical electrodes were placed on the brow bone and eye socket of the left eye. The EEG signals were amplified by a SynAmps amplifier and Curry 7.0 software and filtered with a 0.1-100 Hz bandpass filter. The resistance of all electrodes was kept below 10 kΩ. The EEG signals were recorded using the right mastoid (A2) as the reference electrode with a sampling rate of 1000 Hz.

### 4.4 Data analysis

#### 4.4.1 Risk preference consistency

Risk preference was initially defined based on the difference between the proportion of risky choices made by a participant and those a rational individual would make ^5,49^. A ‘rational individual’ is described as someone who chooses the risky option when its expected value (amount × probability) exceeds that of the safe option (¥10). A positive value indicates a risk-seeking preference, while a negative value suggests risk-aversion.

Next, the consistency of risk preference was defined as the degree to which an individual’s actual choices matched their initial risk preference. This was quantified using the formula: Risk Preference Consistency = Risk Preference × Sign. Here, ‘Sign’ equals 1 for risk-seeking individuals and -1 for risk-averse individuals. Therefore, more negative value indicated that more inconsistent choices were made, reflecting a greater degree to which actual risk preferences contrary to initial risk preferences.

Pre-observation risk preference consistency was measured based on choices in the first round of the task. Post-observation risk preference consistency was calculated as the average of the risk preference consistency observed in the fourth and seventh rounds.

#### 2.4.2 Hierarchical drift-diffusion model (HDDM)

DDM is characterized by several key parameters (see Figure 2.C). Drift Rate (*v*): this measures the speed of evidence accumulation, which can be influenced by the attributes of the options. A more positive or negative value indicates a preference for the upper or lower boundary, respectively (in this study lower boundary representing options increased after observation). Decision Threshold (*a*): this parameter represents the distance between the two boundaries, indicating the amount of evidence needed before a decision is made. Starting point (*z*): this indicates an initial bias towards one of the boundaries at the beginning of the drift process. A larger or smaller value indicates a greater choice bias towards the upper or lower boundary option before any evidence is accumulated. The Bayesian hierarchical drift diffusion model (HDDM) was utilized to assess trial-specific parametric influences on the underlying decision processes ^50^. This model employed a Bayesian estimation approach to calculate the combined posterior distribution of parameters based on observed decision data, including reaction times and choices. The study examined potential changes in the weights of amount, probability or the expect (amount × probability) on drift rate, the decision starting point and the decision threshold, resulting in 24 competing models in each group (see Figure 2.C, detailed in Table S1). The Markov chain Monte Carlo (MCMC) sampling method was employed to approximate the posterior distribution of parameters, generating 12,000 samples while discarding the first 1,000 as burn-in. Reaction times (RT) shorter than 0.3 seconds were excluded from the analysis. To confirm model convergence, R-hat (Gelman-Rubin) convergence statistics were computed, with no R-hat statistics exceeding 1.1, indicating good convergence ^18,51,52^. The Deviance Information Criterion (DIC) was applied for hierarchical model comparison, considering a difference greater than 10 significant ^18,53^. The parameters from the winning model underwent posterior predictive checks to assess whether the model could accurately reproduce key patterns observed in the data.

#### 4.4.3 EEG pre-processing and Time-frequency analysis

EEG data were pre-processed using EEGLAB v13.0.1 and MATLAB R2016b (The Mathworks, Natick, MA). Offline EEG data were down-sampled to 250 Hz and re-referenced to the left mastoids. Data were filtered with a 1–30 Hz bandpass filter via a basic finite impulse response (FIR) filter. Continuous EEG was epoched from -600 to 2000 ms (650 sampling points) for the stimulus marker in the decision phase. Independent Components Analysis (ICA) was used across all epochs to remove ocular artifacts. An epoch would be automatically rejected whenever the voltage reached more than 100 *μ*V.

The processed EEG data were converted into time-frequency domain data using continuous wavelet transform (CWT) in Letswave software (https://www.letswave.org/) ^54^. The values of central frequency (*ω*) and limit (*σ*) were set to 5 and 0.15, respectively, in the continuous wavelet transform. The range of time-frequency representation (TFR) was 1–30 Hz, and the step was 0.58 Hz. The TFRs of a single trial would be obtained after wavelet transform. Then, the spectral perturbation in TFRs was calculated to obtain energy regulation of EEG rhythms and it was defined as: *ER*_t,f_% = [*A*_t,f_ – *R*_t,f_]/*R*_f_ . *A*_tf_ was the energy of single-trial oscillation at a given time *t* and frequency *f* and *R_f_* was the mean of single-trial energy in the baseline time window. -550 to -150 ms before the decision phase was selected as the baseline time window. Considering that previous studies have reported the neural activities in the fronto-parietal regions, we primarily focused on Fz, F3, F4, FCZ, FC3, FC4, CZ, C3, C4, CPZ, CP3 and CP4 for theta (4–7 Hz) and beta (15–30 Hz).

#### 4.4.4 Multivariable analysis for the neural activities associated with DDM parameters

We first conducted inter-subject representational similarity analysis (IS-RSA) to identify the neural activities linked to changed DDM parameters. First, for parameters altered by social influence (such as the possible starting point), we calculated the change in each group (post-observation minus pre-observation). Then, we computed the Euclidean distance between participants based on these change values to construct a parameter-related representational dissimilarity matrix (RDM). Next, for each of the 650 sampling points of neural data, we also calculated the change in each group (post-observation minus pre-observation) and averaged all TFRs within the theta or beta band, resulting in a 1×12 vector for each participant. Based on the vector, we computed the Euclidean distance between participants in each group to obtain theta- and beta-related RDM at each sampling point. All distances were rescaled to the range [0, 1]. Furthermore, at each sampling point, we used a general linear model (GLM) to compute, the prediction of the interaction between each neural RDM and group (dummy code: Con group = 1; Incon group = 2) on the parameter RDM (RDM_parameter_ ∼ *β*1 × RDM_theta_ × group + *β*2 × RDM_beta_ × group). Multiple comparisons were corrected using Family Wise Error (FWE) method. The significant interaction indicated that the prediction of certain neural RDM on the parameter-related RDM differed between groups. For sampling points with significant interaction effects, we used GLM to estimate the prediction of the parameter-related RDM by the neural RDMs in each group to interpret the simple effects. We also explored how social influence shaped (increased or decreased) theta and beta activity. To address this, we averaged the theta and beta activity corresponding to the RSA-significant sampling points in each group. Then linear mixed-effects model where participant was incorporated as a random intercept to examine the interaction effect between group (dummy code: Con group = 1; Incon group = 2) and time (dummy code: pre-observation = 1; post-observation = 2) on theta and beta activity, respectively.

Finally, to further support that the identified neural activity underlies social influence, we employed a support vector machine (SVM) approach to examine whether the neural activity could effectively distinguish between the two groups. Specifically, for each participant, we extracted both theta and beta activity at time points showing significant interaction effects and concatenated them into a 3D matrix (participants × electrodes × sampling points). We then averaged across the sampling points, resulting in a 59 (participants) × 12 (electrodes) matrix, which was subsequently rescaled to the range [0, 1]. We labeled the Incon group as 1 and the Con group as -1, and performed classification using a linear SVM implemented with the LIBSVM toolbox ^55^. The linear kernel was selected to reduce the risk of overfitting and to allow straightforward interpretation of feature weights. The SVM’s regularization parameter *C*, which controls the balance between training accuracy and model generalization, was set to its default value of 1. The null distribution of classification accuracy was generated through 5,000 permutation tests. The *p*-value was defined as the proportion of permuted accuracies exceeding the actual accuracy. *P* < 0.05 was considered statistically significant.

## Notes

**Funding disclosure statement:** This work was supported by grants from the Brain Science and Brain-like Intelligence Technology—National Science and Technology Major Project (2025ZD0215701) and National Natural Science Foundation of China (32071082 and 71942001).

### Competing Interest Statement

The authors have declared no competing interest.

